# Regulation of Ordinal DNA Translocation Cycle in Bacteriophage Φ29 through Trans-Subunit Interactions

**DOI:** 10.1101/2025.01.29.635585

**Authors:** Rokas Dargis, Joshua Pajak, Pavan Ariyawansa, Marc C. Morais, Paul J. Jardine, Gaurav Arya

## Abstract

Certain viruses such as tailed bacteriophages and herpes simplex virus package double-stranded DNA into empty procapsids via powerful, ring-shaped molecular motors. High resolution structures and force measurements on the DNA packaging motor of bacteriophage Φ29 revealed that its five ATPase subunits coordinate ATP hydrolysis with each other to maintain the proper cyclic sequence of DNA translocation steps about the ring. Here, we explore how the Φ29 motor regulates translocation by timing key events, namely ATP binding/hydrolysis and DNA gripping, through trans-subunit interactions. We used subunit dimers bound to DNA as our model system, a minimal system that still captures the conformation and trans-subunit interactions of the full pentametric motor complex. Molecular dynamics simulations of all-ATP and mixed ATP-ADP dimers revealed that the nucleotide occupancy of one subunit strongly affects the ability to hydrolyze ATP in the adjacent subunit by altering the free energy landscape of its catalytic glutamate approaching the gamma phosphate of ATP. Specifically, one ATP-bound subunit donates residues in trans that sterically block the neighboring subunit’s catalytic glutamate. This steric hindrance is resolved when the first subunit hydrolyzes ATP and is ADP-bound. This obstructive mechanism is supported by functional mutagenesis and appears to be conserved across several Φ29 relatives. Mutual information analysis of our simulations revealed intersubunit signaling pathways, via the trans-acting obstructive residues, that allow for sensing and communication between the binding pockets of adjacent subunits. This work shows that the sequential order of DNA translocation events amongst subunits is preserved through novel trans-subunit interactions and pathways.

## Introduction

To package their genomes into procapsids, double-stranded DNA (dsDNA) viruses have to be able to generate large forces to translocate DNA against resistance arising from electrostatic repulsion, bending, and entropic loss of tightly-packaged DNA [2, 3]. Such viruses utilize a powerful, ring-shaped motor that hydrolyzes ATP to generate the required mechanical force. These ATPase motors belong to the ASCE (additional strand conserved glutamate) family and are characterized by conserved Walker A and Walker B motifs, which are involved in nucleotide binding, Mg^2+^ binding, and catalyzing hydrolysis [1, 4]. Thus, understanding how these motors function would elucidate not only viral DNA packaging, but also mechanisms of other processes, such as genome segregation, cell division, and protein degradation which also utilize ASCE motors for substrate translocation [4]. Furthermore, insights into viral packaging motors could inform the development of novel antiviral therapies since many disease-causing viruses, such as herpes, also utilize a similar packaging system [5].

The Φ29 packaging motor is an excellent model system to study genome packaging motors due to the availability of an efficient *in vitro* packaging system, which facilitate single-molecule investigations of DNA packaging [6, 7, 8, 9], and the only near-atomic resolution structure of an intact viral packaging motor complex [1], which provides unprecedented structural insight. Additionally, this structure can be used as a starting model for performing molecular dynamics (MD) simulations [10, 11] for studying the dynamic aspects of the motor. Bacteriophage Φ29 motor ring consists of five identical subunits of gene product 16 (Fig. 1 A), each composed of three main domains: a vestigial nuclease C-terminal domain (CTD) [11, 12] that attaches to a cone-shaped complex (connector) located at pro-capsid portal [13], an N-terminal ATPase domain (NTD), and a linker domain that connects the CTD and NTD. Encompassed within the linker domain is the “lid” structure that partially engages with the neighboring subunit (Fig. 1 B).

**Figure 1.**
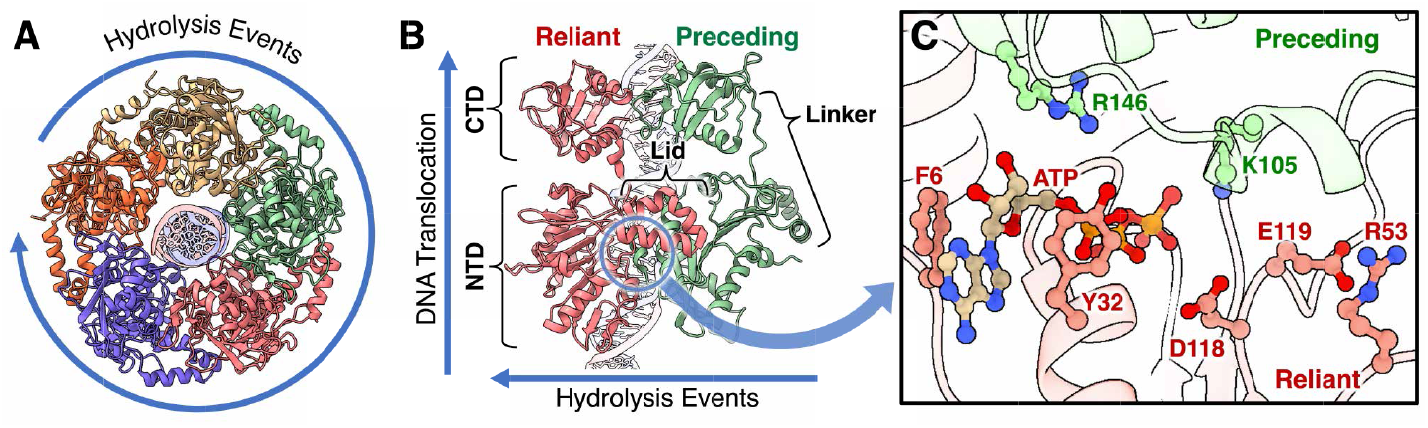
Structure of Φ29 gp16 motor with DNA. (A) Axial view of the pentameric motor, where the direction of DNA translocation is out of the page towards the reader. The outer arrow signifies the cycle of hydrolysis events. Subunits are colored differently and DNA can be seen in the center channel. (B) Side view of the second (green) and third (pink) subunits in the cycle. Based on the order of hydrolysis, the green subunit is labeled as the preceding subunit and the pink one is the reliant subunit. Each subunit consists of a NTD and CTD connected by a linker domain. The lid structure of the reliant subunit engages with the NTD of the preceding subunit. (C) Zoomed in view of the reliant subunit’s binding pocket, located at the interface with the preceding subunit. Key residues and ATP are portrayed in the stick and ball format, while the rest of the protein is in the cartoon format and translucent. Both subunits have residues that interact with ATP. E119 is the cis-catalytic glutamate, R53 is the cis-glutamate switch, K105 is the trans-catalytic residue, and R146 is the trans-nucleotide exchange residue. F6 and Y32 help stabilize the ATP. PDB: 7JQQ [1]

Recently, a detailed mechanism for DNA packaging was proposed based on single-molecule, structural, and computational work on Φ29 and related motors [11]. In this mechanism, the motor ring alternates between planar and helical conformations, corresponding to the dwell and burst phases of the packaging cycle observed experimentally. The dwell phase begins with the motor in the planar conformation. One by one, in each subunit, ATP replaces ADP in the binding pocket [9], causing the NTD of that subunit to reach down (away from CTD; see Fig. 1 B) and grip the dsDNA. Each consecutive subunit will need to extend further to reach the DNA backbone, imposing increasingly unfavorable rotations on the lid structure. The end of the dwell phase is characterized by all subunits being ATP bound and gripping dsDNA, leading to a helical conformation of the NTDs, which roughly matches the helicity of the DNA backbone. This configuration is now ready to enter the burst phase, wherein ATP hydrolysis proceeds ordinally around the motor. When a subunit hydrolyzes ATP and releases its grip on DNA, the neighboring ATP-bound subunit is no longer constrained to the extended conformation and its lid structure rotates back to release the helical strain. This allows the neighboring subunit’s NTD to become coplanar with the hydrolyzing subunit and brings the NTD ring closer to planarity, translocating DNA in the process. Each translocating step is roughly 2.5 basepairs (bp), and the full burst phase translocates the dsDNA by one helical turn [7]. At the end of the burst phase, the motor has returned to a planar conformation and is now fully ADP bound, ready to re-enter the dwell phase.

Single-molecule force spectroscopy (SMFS) experiments were the first to show that the hydrolysis of ATP proceeds cyclically about the subunit ring, suggesting a high degree of inter-subunit coordination [9, 7]. In particular, a subunit will not proceed with ATP hydrolysis until the subunit prior to it in the hydrolysis cycle has already hydrolyzed ATP. This coordination must involve the interface of the two subunits, which is where the ATP binding pocket is located. While ATP binds to only one subunit, its hydrolysis mechanism *relies* on residues of the *preceding* subunit as well; hereon adjacent subunits are referred to as “reliant” and “preceding” (Fig. 1 C). Since the coordination of hydrolysis between subunits is likely linked to their relative positions within the planar/helical ring, the mechanism by which the binding pockets communicate with each other to maintain ordinality is unclear.

Here we use all-atom MD simulations, free energy calculations, and mutual information analysis, supplemented by experimental mutagenesis, to show that the glutamate switch mechanism—previously thought to regulate hydrolysis and DNA gripping *in cis* within individual subunits [14, 15]—is further regulated *in trans* by a neighboring subunit.

We delineate how this regulation, which occurs through an obstructive motif at the subunit interface, couples the nucleotide state of the preceding subunit to the hydrolysis event in the reliant subunit. Broadly, our results suggest that regulation mechanisms in translocation motors likely involve closely interlinked cis- and trans-mechanisms[15, 1].

## Results

### Minimal Model for Studying Trans-Interactions

The full pentameric Φ29 ATPase motor, bound to dsDNA, is rather large for MD simulations; due to the symmetry of the motor, not all parts of the motor are required to analyze trans-interactions between two subunits. Therefore, our first objective was to determine the minimal composition of the motor that could still accurately capture trans-subunit interactions. We started off by carrying out equilibrium simulations of a single ATP-bound subunit (Fig. 2 A) and monitoring its structural stability through root mean square deviation (RMSD) of the protein’s backbone. The simulations found that isolating a monomer leaves it unstable (RMSD of over 10 Å; Fig. 2 B, blue line) due to the loss of structural support provided by adjacent subunits and dsDNA. The linker domain, and in particular the lid structure, which no longer has a neighbor to engage with, becomes unstable and leads to large movements of the CTD and NTD relative to each other that would not be expected of subunits that are part of the pentamer (SI Appendix, Fig. S1 A).

**Figure 2.**
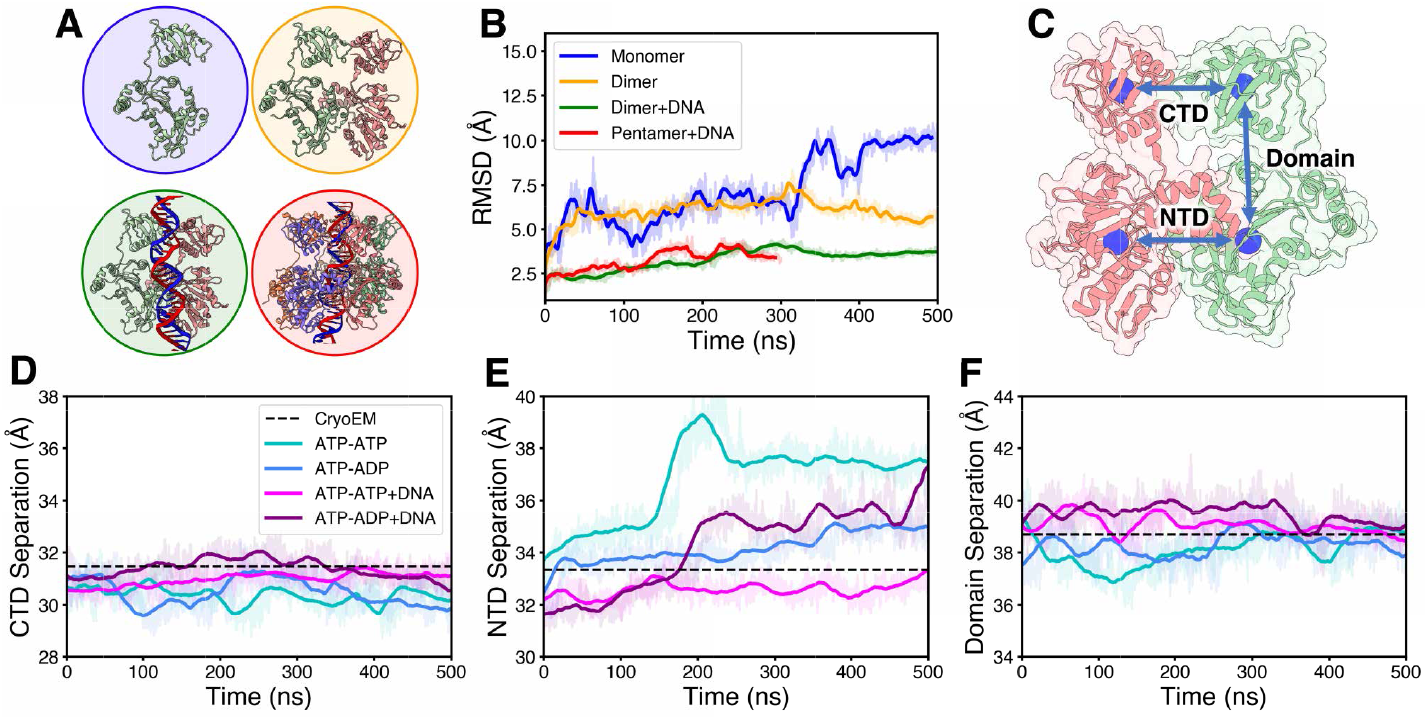
Analysis of various structure compositions of the Φ29 motor. (A) Monomer (blue), dimer (orange), dimer with DNA (green), and pentamer with DNA (red). (B) RMSD of the third subunit in the pentameric motor with DNA compared to the same subunit in monomer, dimer, and dimer with DNA models. (C) Reference dimer showing how CTD, NTD, and domain (CTD-NTD) separation distances (arrows) are calculated using center of masses (circles). CTD separation (E), NTD separation (D), and domain separation of the preceding subunit (F) of ATP-ATP and ATP-ADP dimers with and without DNA. In panels B, D, E, and F, shaded lines represent frame-by-frame values while the solid lines are the moving average. PDB: 7JQQ [1]

We then carried out simulations of a system that consisted of two neighboring ATP-bound subunits from the cryo-EM structure [1] (Fig. 2 A). The dimer system allows the reliant subunit to have a stable lid structure through contact with the neighboring subunit. While this system was found to be more stable than the monomer and the reliant subunit equilibrated at an RMSD of 6 Å (Fig. 2 B, orange line), the subunits began drifting apart at the NTD interface (SI Appendix, Fig. S1 B).

To investigate if the presence of dsDNA next to subunits prevents the observed drift, we next simulated the same dimer system, but bound to dsDNA as found in the cryo-EM structure (Fig. 1 A). Now, the reliant subunit equilibrated at a lower RMSD of 4 Å (Fig. 2 A, green line), indicating increased stability. Given that a resistive force of 57 pN is required to stall the motor [6] and, as described in the helical-planar ratchet mechanism [11], the conformation of the motor is dictated by the gripping and release of dsDNA, we attribute the increased stability to the strong interaction between the dsDNA and a subunit’s DNA gripping residues.

In fact, this stability is comparable to that of the simulated ATP-bound pentamer with dsDNA (Fig. 2 A, B, red line). Having shown that our isolated DNA-bound dimer system captures the relevant properties of a representative dimer in the pentamer before hydrolysis (both the reliant and preceding subunit are ATP bound), we next tested whether this system also captured the post hydrolysis state. Therefore, we also simulated a DNA-bound ATP-ADP dimer (preceding subunit now ADP bound), and compared relevant distances between their domains (CTD-CTD, NTD-NTD, and CTD-NTD) to those in the cryo-EM structure (Fig. 2C-F).

While the CTD-CTD and CTD-NTD distances remained consistent with values in the cryo-EM structure regardless of the preceding subunit’s nucleotide occupancy (Fig. 2 D, F), we find that the DNA-bound ATP-ADP dimer exhibits larger NTD-NTD distances (Fig. 2 E). This difference can be explained by SMFS experiments, which show that ATP-bound subunits grip dsDNA tighter than ADP-bound subunits; therefore, when DNA is present, the NTD domains are held more closely together in the ATP-ATP system compared to the ATP-ADP system. Indeed, we find that the removal of DNA from the dimer complex led to unstable NTD-NTD distances in both the ATP-ATP and ATP-ADP configuration; the CTD distances remained unaffected (Fig. 2 C-F).

Taken together, our results suggest that the DNA bound dimer system is a computationally optimal model for studying trans-subunit interactions because it captures both the conformation of and the interactions between subunits of the pentameric motor. Henceforth, all remaining analysis will be carried out using this model system, ATP-ATP for pre-hydrolysis and ATP-ADP for post-hydrolysis states.

### Trans-Dependence of Glutamate Switch Mechanism

The glutamate switch is a crucial mechanism for regulating ATP hydrolysis in cis. This mechanism involves three components: the catalytic glutamate (E119), the switch (R53), and the gamma-phosphate of ATP (P_*γ*_) [15]. In the “inactive” state, the catalytic glutamate is interacting with the switch and is prevented from reaching P_*γ*_ . In the “active” state, the catalytic glutamate dissociates from the switch and approaches P_*γ*_, allowing it to catalyze ATP hydrolysis by deprotonating a water molecule that will perform a nucleophilic attack on P_*γ*_ [4] (Fig. 3 B). Previous free energy calculations showed that the state of the glutamate switch is dependent on the nucleotide state of the subunit: an ATP-bound monomer favors the active state, whereas an ADP-bound monomer favors the inactive state and has a wider and higher free energy barrier separating the two states [15].

**Figure 3.**
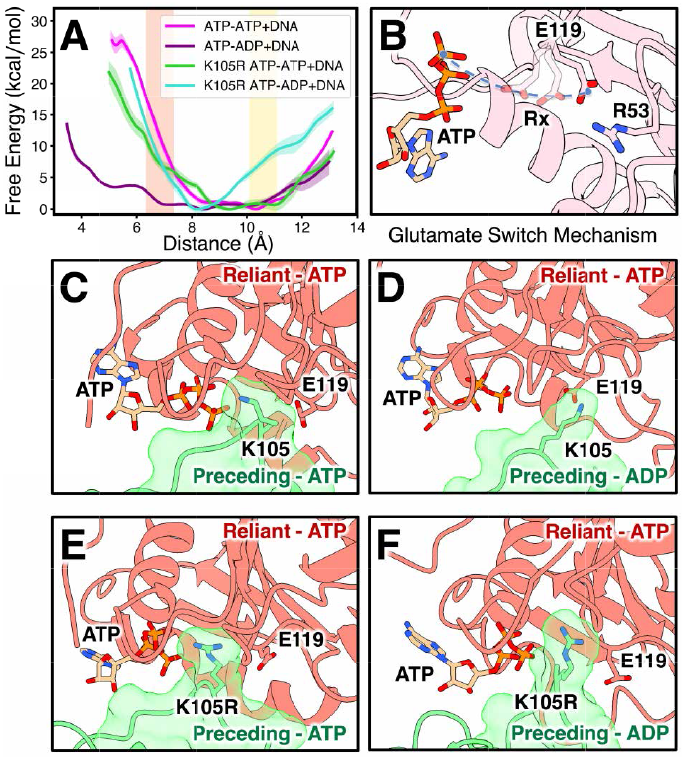
Effect of the obstructive motif on the glutamate switch mechanism. (A) Free energy landscape of the reliant subunit’s catalytic glutamate (E119) approaching the P_γ_ in ATP-ATP and ATP-ADP DNA dimers (pink and purple lines) with WT obstructive lysine (K105) and mutant K105R (green and cyan lines). Shaded zones around the lines represent error bars. The active and inactive states of the glutamate switch mechanism are highlighted in red and yellow. (B) Visualization of the glutamate switch mechanism. The catalytic glutamate is shown in the “inactive” state (opaque E119) when interacting with the glutamate switch (R53) and transitioning towards the “active” state (translucent E119) when approaching P_γ_ of ATP. The reaction coordinate (blue circles and dotted line) of free energy calculations is the distance between the COM of the oxygen in the active group of E119 and P_γ_ . Visualization of the obstructive lysine in the WT ATP-ATP DNA (C), WT ATP-ADP DNA (D), K105R mutated ATP-ATP DNA (E), and K105R mutated ATP-ADP DNA dimers (F).

To test if the state of the glutamate switch (active *vs*. inactive) in the reliant subunit also depended on the nucleotide occupancy of the preceding subunit, we calculated free energy, using umbrella sampling simulations, as a function of the distance between the catalytic glutamate and P_*γ*_ of ATP in the reliant subunit of our pre-and post-hydrolysis dimer model system. The ATP-ATP dimer exhibited an energy minimum near the inactive state and a free energy increase of 10 kcal/mol to reach the active state, whereas the ATP-ADP dsDNA dimer had a relatively flat free energy landscape, with a difference of *<* 5 kcal/mol between the two states (Fig. 3 A). This suggests that the reliant subunit cannot hydrolyze ATP until the preceding subunit has first hydrolyzed its ATP, which is consistent with our helical-planar ratchet mechanism which suggested that the adjacent subunits help maintain the ordinality of ATP hydrolysis through trans-interactions [11].

### Regulation through Steric Hindrance

To identify the molecular basis for the difference in the free energy landscapes, we compared the subunit interface of ATP-ATP and ATP-ADP dimers at the binding pocket of the reliant subunit. From our equilibrium simulated trajectories, we observed that in the ATP-ATP dimer, residues numbered 100 to 108 of the preceding subunit form a helical structure and position themselves between the reliant subunit’s catalytic glutamate and ATP, preventing the catalytic glutamate to transition from the inactive to the active state (Fig. 3 C, SI Appendix, Fig. S2). In contrast, this “obstructive motif” moves out of the way in the ATP-ADP dimer (Fig. 3 D, SI Appendix, Fig. S2, S4), facilitating the transition to the active state and thereby ATP hydrolysis. Thus, this helical obstruction contributes to the additional energy required by the catalytic glutamate to transition to the active state in the ATP-ATP dimer compared to its ATP-ADP counterpart (Fig. 3 A).

Interestingly, lysine 105 (K105) is part of the obstructive motif and is the primary residue that positions itself between the catalytic glutamate and ATP. This residue has previously been proposed to be a trans-acting catalytic residue that aids in the cleaving of P_*γ*_ during hydrolysis [1], akin to the highly conserved “arginine finger” found in many related ATPase motors [16]. In simulations where the preceding subunit is ATP bound, K105 of the preceding subunit is seen to be consistently interacting with the P_*γ*_ of the ATP bound to the reliant subunit (within 3 to 5 Å of each other). However, when the preceding subunit is ADP bound, its K105 becomes more promiscuous and can be found interacting not only P_*γ*_, but also the catalytic glutamate of the reliant subunit, and sometimes neither (up to 13 Å away from P_*γ*_). This change in promiscuity appears correlated to the *χ*_1_ angle of K105, which was found to be more stable in the ATP-ATP dimer versus the ATP-ADP dimer (SI Appendix, Fig. S3).

Tying these results together, we propose that the preceding subunit can regulate the hydrolysis of the reliant subunit via the preceding subunit’s obstructive motif, whose position is tied to the preceding subunit’s nucleotide state despite being located over 40 Å away.

### Allosteric Pathways Connecting Key Residues

Interactions in macromolecular complexes can be regulated through long-range allosteric communication, where conformational changes (ordered or disordered) in one part of the protein induce changes in a different part of the protein or another subunit [17]. We next probed for an allosteric link between the binding pockets of adjacent subunits to elucidate how the information of the preceding subunit’s nucleotide state is transmitted to the reliant subunit. To this end, we applied the Correlation of All Rotameric and Dynamical States (CARDS) method [17] on our equilibrium simulation trajectories to calculate the mutual information (MI) from the preceding subunit’s catalytic glutamate to all dimer residues. The catalytic glutamate was chosen because its state (active *vs*. inactive) is representative of the subunit’s nucleotide state (ATP *vs*. ADP).

Our analysis revealed that the target catalytic glutamate of the preceding subunit is not directly linked to the catalytic glutamate of the reliant subunit, but rather to residues of the preceding subunit that act in trans on the reliant subunit’s binding pocket (Fig. 4 B), namely K105, which we just showed was part of the obstructive motif, and R146, which has previously been implicated in facilitating nucleotide exchange in trans [1]. A second MI calculation, this time using the preceding subunit’s K105 as the target residue, showed a correlation to the reliant subunit’s catalytic glutamate and R146, and the preceding subunit’s K124 (Fig. 4 B), a proposed DNA gripping residue [1, 18]. Combining the two MI calculations, we infer that the preceding subunit’s catalytic glutamate is indirectly linked to the reliant subunit’s catalytic glutamate via the preceding subunit’s K105, which further supports its role as a regulating mechanism of ATP hydrolysis. We also find that K105 acts as a signal hub based on its connections to additional key residues, providing a mechanism for the motor to coordinate DNA gripping and nucleotide exchange based on the current nucleotide state of the subunit and its neighbors.

**Figure 4.**
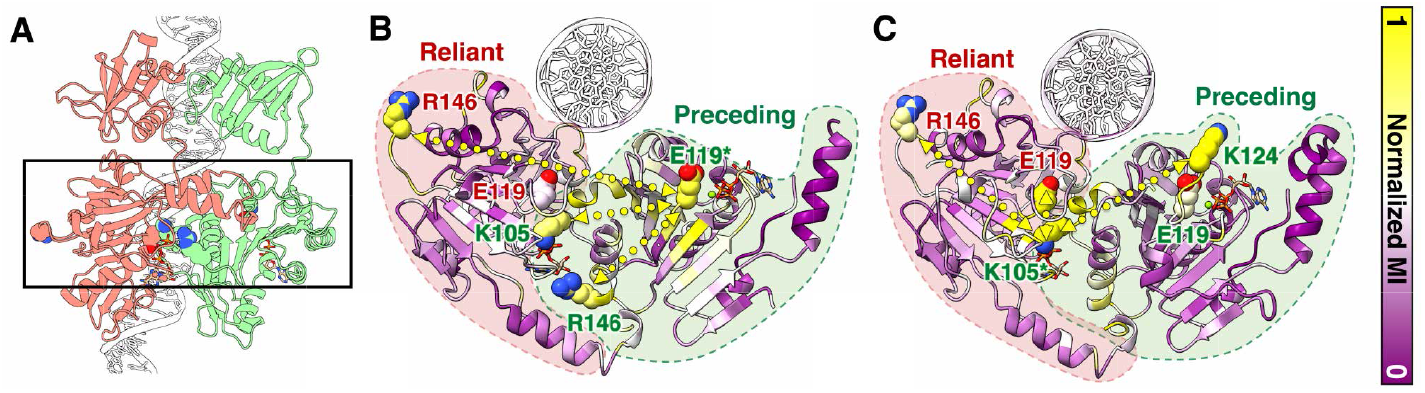
Normalized mutual information calculations across various ATP-ATP DNA dimer simulations. (A) Side view of the dimer model, highlighting the focus region (box) of the next two panels. (B) MI of preceding subunit’s catalytic glutamate (E119). (C) MI of the preceding subunit’s obstructive lysine (K105). The target residue is marked with an asterisk, and the color of the residue label corresponds to which subunit the residue belongs. Dotted yellow arrows denote suggested allosteric pathways. Both panels utilize the same color bar for MI but have been individually normalized with outliers removed.

### Mutagenic and Phylogenetic Evidence

To experimentally demonstrate that K105 regulates ATP hydrolysis through steric hindrance and not just trans-catalytic interactions, we performed poisoning experiments where the ATPase activity of Φ29 motors with varying ratios of wild-type (WT) and mutated subunits were tracked. In the first poisoning experiment, the mutant was a replacement of K105 with arginine (K105R) and was selected because both lysine and arginine have a similar positively charged active group, but arginine has a slightly longer side chain. If the role of K105 is purely trans-catalytic, we expect that K105R would have comparable ATPase activity based on similar systems, such as the F_1_-ATPase [19], which utilize an arginine in that role. However, if the obstructive motif functions as proposed here, the slightly bulkier side chain would make it harder for the obstruction to move out of the way of the catalytic glutamate, and ATPase activity would be expected to decrease with higher ratios of K105R subunits.

The poisoning experiments show that increasing the proportion of K105R over WT subunits decreases the ATPase activity (Fig. 5 A), which is in line with the proposed obstructive regulatory role of K105. To confirm that the mutant negatively impacts the obstructive mechanism, we repeated the free energy calculations, in which the preceding subunit now had the K105R mutation (Fig. 3 A). The free energy landscape did not change significantly between the native and mutated ATP-ATP dimers. However, the mutated ATP-ADP dimer incurs a larger energy penalty to transition from inactive to active compared to the native ATP-ADP dimer. This is further supported by equilibrium simulations of the mutated dimers which showed that K105R has lower rotational freedom (SI Appendix, Fig. S3) and could not move out of the way to unblock the catalytic glutamate within the simulation timescale (Fig. 3 E, F). Additionally, MI calculations revealed a decrease in correlation between the catalytic glutamate and K105R (SI Appendix, Fig. S6). Together, these *in silico* analyses provide a molecular basis for the decreased ATP activity in the K105R mutant.

**Figure 5.**
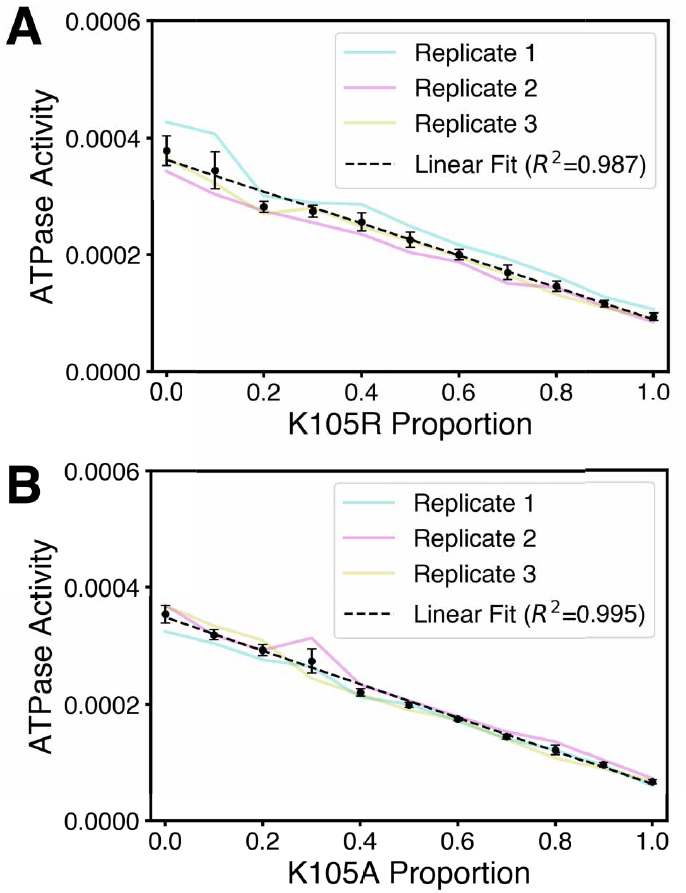
Non-packaging ATPase activity of K105R (A) and K105A (B) poisoning experiments. Proportion signifies the average fraction of mutant subunits in the assembled pentamers.

The next poisoning experiment involved an alanine mutant of K105 (K105A). Alanine was chosen due to its small and inert side chain, which is expected to decrease the size of the obstruction. Similar to K105R, the ATPase activity decreased as the proportion of K105A mutant subunits increased (Fig. 5 B). However, since this mutant eliminates both the obstructive element by reducing the size of the side chain and the catalytic function by removing the active group, determining which of the two roles led to the decrease of ATPase activity becomes difficult. We believe that the overall decrease in ATPase activity with the K105A mutant may be a compounding effect: the removal of the active group leads to a large decrease in activity and the reduction of the obstruction leads to a smaller increase in activity.

Lastly, we found evidence of the obstructive motif mechanism in several phylogenetic relatives of Φ29: GA-1, SF5, B103, and Φ28 [20, 11] [SF5 sequence pending]. Since we do not have atomic resolution structures of these relatives, besides Φ28, AlphaFold [21] was used to generate their monomeric structures. The NTDs of two of these monomers were then superimposed onto the NTDs of a Φ29 dimer, allowing us to create a reasonable model of their subunit-subunit interface. Notably, dimers of these relatives were found to contain a helical structure with a lysine that is properly positioned to obstruct the catalytic glutamate in a manner very similar to Φ29 (Fig. 6). Even if their packaging mechanism may not be the same as that of Φ29, the fact that all of these systems have an obstructive helix-like structure leads us to believe that it is important and universal enough to be conserved.

**Figure 6.**
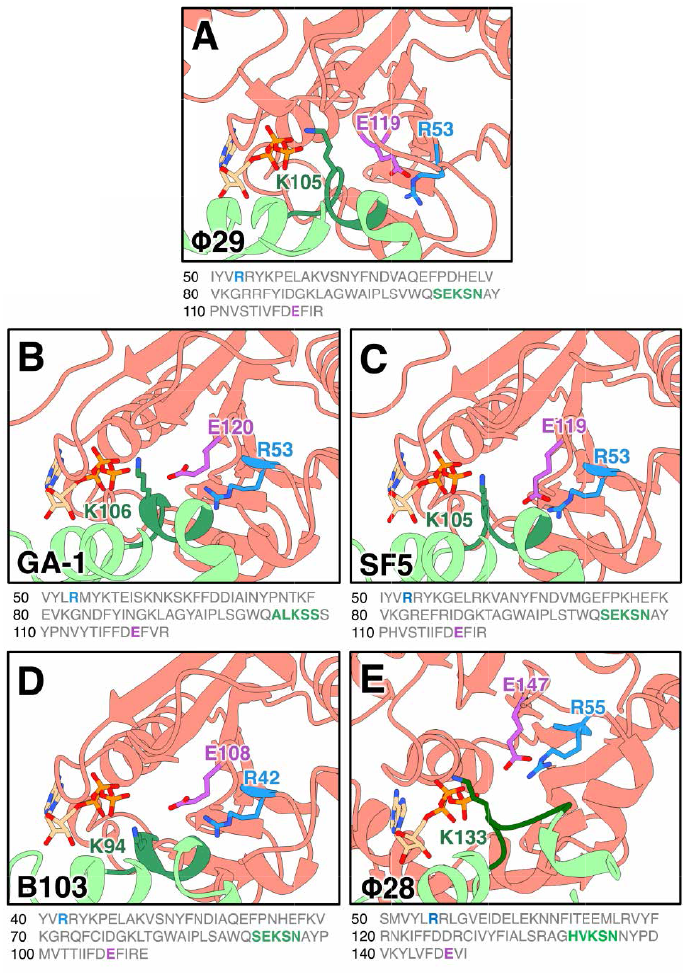
Visualization of the binding pocket and relevant residues of Φ29 motor and its relatives. (A) Obstructive motif of Φ29 dimer (preceding subunit is green and reliant is red) as seen in the cryo-EM reconstruction. This obstructive motif is also seen in the dimer reconstruction of Φ29 relatives: GA-1 (B), SF5 (C), B103 (D), and Φ28 (E). Corresponding sequences highlight the arginine that acts as the glutamate switch (blue), the obstructive motif (green) and the catalytic glutamate (purple).

## Discussion

Our current understanding of viral DNA packaging by the Φ29 ATPase motor is that it operates in two phases [7]: in the burst phase, the motor transitions from a helical to planar conformation as subunits hydrolyze ATP, thereby generating the required force to translocate DNA; in the dwell phase, the motor returns to the helical conformation as ATP replaces ADP in each subunit [1]. Remarkably, both hydrolysis and nucleotide exchange occur cyclically and ordinally about the motor subunits, suggesting a high degree of coordination. While it is evident that trans-subunit interactions must be responsible for maintaining ordinality, the underlying molecular mechanisms remain unknown.

In this work, we propose a novel mechanism that coordinates ATP hydrolysis between neighboring subunits in the burst phase. Specifically, based on its nucleotide occupancy, a subunit is able to sterically prevent the next subunit from hydrolyzing out of turn. MD simulations using our DNA-bound dimer model, which we showed is sufficient to study communication between adjacent subunits, revealed that the coordination mechanism consists of two core residues: a catalytic glutamate (E119) and an obstructive lysine (K105). When the preceding subunit is still ATP bound, its obstructive lysine positions itself between the catalytic glutamate and P_*γ*_ of the ATP bound to the reliant subunit (next to hydrolyze), thus hindering its hydrolysis (Fig. 7 B top). Hydrolysis in the preceding subunit causes the obstructive lysine to retract as a result of signaling pathways that connect it to the same subunit’s catalytic glutamate (Fig. 7 A, cyan arrow), as captured by our MI calculations. This retraction allows the catalytic glutamate to adopt the active pose and participate in hydrolysis (Fig. 7 B, bottom). Using free energy calculations, we quantified the energetics associated with the movement and removal of the obstruction using both native and lysine-mutated subunits.

**Figure 7.**
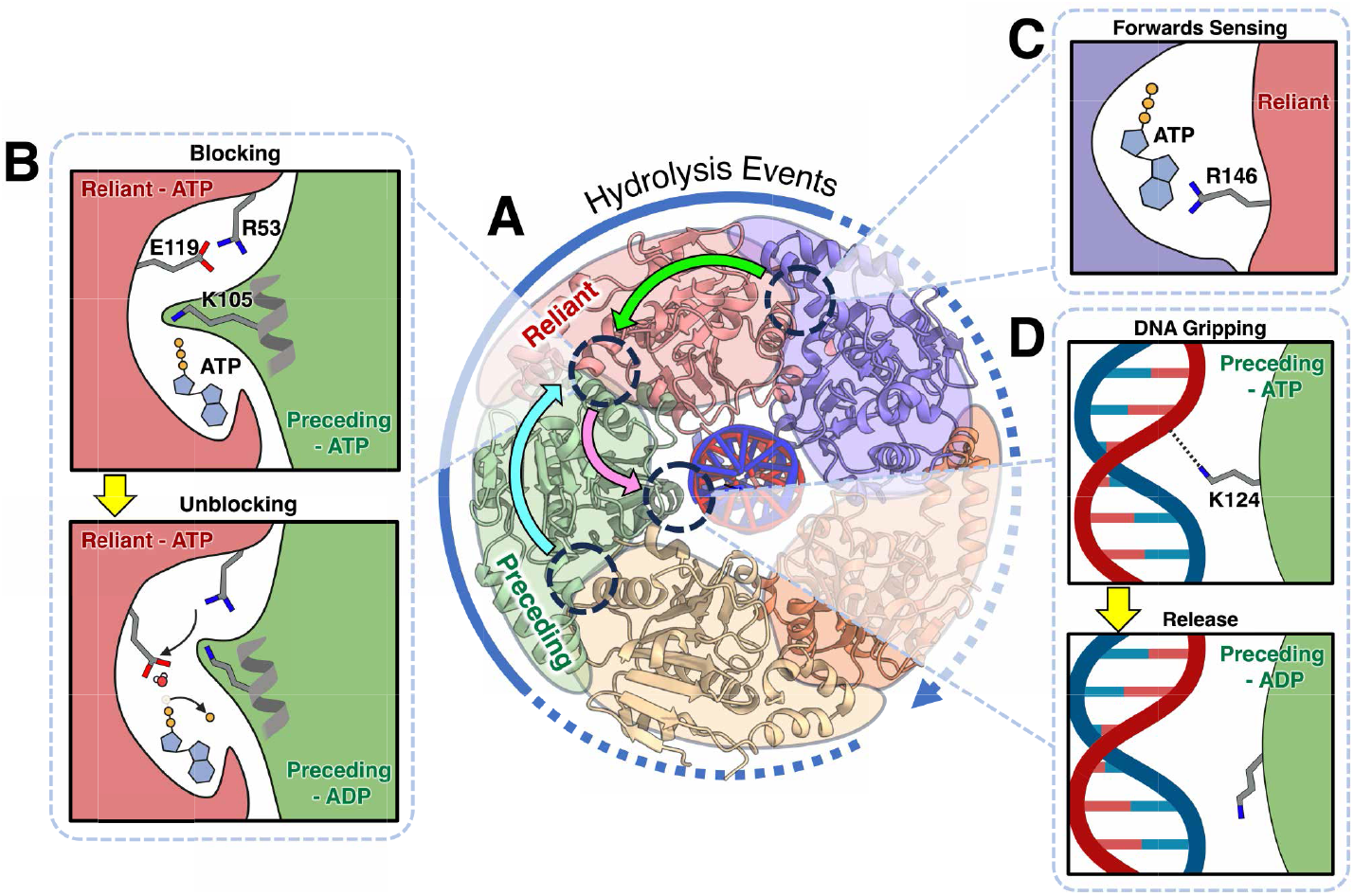
Proposed mechanism by which the Φ29 motor maintains ordinality in the burst phase of the DNA translocation. A) Identified communications pathways (arrows) between key regions (dotted circles) of adjacent subunits. Spotlights lead to a detailed view of the interactions found in the key regions. The mechanism starts in the binding pocket of the preceding subunit (green), where a hydrolysis event leads to a conformation change in the preceding subunit’s catalytic glutamate. This is communicated to the preceding subunit’s obstructive motif (cyan arrow), which is blocking the reliant subunit’s catalytic glutamate before hydrolysis in the preceding subunit (B, top) but moves out of the way after hydrolysis (B, bottom). The blocking and unblocking effect by K105 is also connected to two additional interactions: (C) proposed forward sensing role via reliant subunit’s R146 (green arrow) which ensures the subunit downstream of the reliant subunit is ATP bound; and (D) gripping and ungripping of DNA via K124 (pink arrow). Coloring scheme and perspective of panel A matches that of Fig. 1, A. The solid part of the dotted “Hydrolysis Events” arrow shows that the stage of the cycle currently involves the green and red subunits.

Our MI analysis revealed two additional allosteric pathways connected to the obstructive lysine. The first pathway connects the lysine to the DNA-gripping residue K124 of the same subunit (Fig. 7 A pink arrow), potentially linking the movement of the obstructive lysine to DNA gripping and release. Prior research established a different allosteric connection, between the catalytic glutamate and DNA gripping residue K56 [15]. Together, these findings suggest that a subunit will release K56 first and then initiate hydrolysis in the next subunit before releasing K124’s grip on DNA (Fig. 7 D), rather than having the hydrolysis event immediately lead to the release of multiple DNA gripping residues. The second pathway connects the preceding subunit’s K105 to the reliant subunit’s R146, which is positioned to interact *in trans* with the adenosine group of the adjacent subunit’s nucleotide (Fig. 7 A green arrow, D). We propose that R146 could play a “sensing” role that checks if the subsequent subunit is ATP bound before starting hydrolysis (Fig. 7 D). This conjecture is in line with the helical-planar ratchet mechanism, where a subunit should only undergo hydrolysis if the subunit before it has hydrolyzed (ADP bound) and the subunit in front of it has not (ATP bound).

While the obstructive mechanism appears to play an important role in maintaining ordinality within the burst phase by utilizing K105 as a checkpoint between key events, the proposed mechanism does not discredit the previously suggested role of K105 as a trans-catalyzing “arginine finger” [1]. We instead suggest that the residue plays two roles: initially aligning itself to hinder the function of the catalytic glutamate, then repositioning itself to unblock the catalytic glutamate and aid in catalyzing hydrolysis. This obstructive mechanism also raises an interesting question regarding the communication between the first and final subunits, which present a unique interface due to the geometry of the motor motif observed in the burst phase. Thus, the question of how these two subunits communicate to initiate the first hydrolysis event of a burst phase still remains open.

For systems that require precise coordination between subunits, having the adjacent subunit directly block another subunit’s action until a condition is met is intuitively very robust. Thus, we propose that our mechanism could be generalized to other viral packaging motors. Indeed, we found evidence of this through structural and sequence comparisons of Φ29 relatives. It is not uncommon to see an overlap of motifs amongst these motors, especially ones that are closely related. The regulation of ATP hydrolysis is present in these motors for a reason, and we expect that deregulating it may negatively impact the overall packaging process even if it speeds up the individual steps.

## Methods

### ATPase assays

ATPase activity of prohead/gp16 motor complexes was determined by measuring production of inorganic phosphate by the EnzChek Phosphate Assay Kit (Life Technologies) as described previously [22]. Briefly, a reaction mixture containing reaction buffer (either kit buffer [50 mM Tris, pH 7.5, 1 mM MgCl_2_, 0.1 mM sodium azide] or TM buffer [25 mM Tris, pH 7.6, 5 mM MgCl_2_]), 0.2 mM of MESG (2-amino-6-mercapto-7-methylpurine riboside) with proheads (4.2 nM) and gp16 (125 nM) in 90 μL was preincubated at room temperature for 10 min in the presence of PNP (purine nucleoside phosphorylase, 0.1 unit). ATP was added to 1 mM to initiate the reaction and production of Pi measured in the spectrophotometer at 360 absorbance for 10 min. For the mutant poisoning assay, the WT and mutant gp16 were mixed in the ratios indicated to yield the 125 nM gp16 concentration.

### Production of proheads, gp16, and gp16 mutants

Proheads were isolated from an infection with a *sus*8.5(900)-*sus*16(300)-*sus*14(1241) mutant (defective in the nonessential head fibers, the packaging ATPase, and the holin, respectively) of *Bacillus subtilis* 12A (nonpermissive host) and purified by sucrose gradient centrifugation as previously described [23].

gp16 was produced by over-expression in *E. coli*. Rosetta cells containing the plasmid carrying SUMO-gp16 were grown overnight in LB medium containing 50 μg/ml kanamycin and 34 μg/ml chloramphenicol at 37°C. The next morning, the culture was diluted 1/100 in LB medium containing 50 μg/ml kanamycin and 34 μg/ml chloramphenicol and grown at 37°C to an optical density at 600 nm of 0.5. Isopropyl-β-d-thiogalactopyranoside was added to 0.3 mM, and the culture was incubated at 37°C for 5 min and then shifted to 18°C for protein expression.

After overnight induction, the cells were collected by centrifugation, concentrated 15-fold in buffer containing 50 mM Tris-HCl, pH 8, 500 mM NaCl, 10% glycerol (vol/vol), and 1 mM tris(2-carboxyethyl)phosphine (TCEP), and lysed by passage through a French press. MgCl_2_ was added to 2.0 mM, DNase was added to 5 μg/ml, and the sample was incubated at room temperature for 15 min. The lysate was clarified by spinning in an SS34 rotor at 10,000 rpm for 40 min at 4°C.

The supernatant was incubated with Talon resin (Clontech) that had been equilibrated in wash buffer (50 mM Tris-HCl, pH 8, 400 mM NaCl, 10% glycerol (vol/vol), 1 mM TCEP at 4°C for 30 min. The resin with SUMO-gp16 bound was then washed with 10 volumes wash buffer. The SUMO-gp16 fusion protein was eluted in 5 volumes elution buffer (50 mM Tris-HCl, pH 8, 400 mM NaCl, 10% glycerol [vol/vol], 100 mM imidazole [pH 8], 1 mM TCEP), and 0.5-ml fractions were collected. The fractions were analyzed by SDS-PAGE to identify peak fractions.

To cleave the SUMO tag and recover native gp16 protein, 100 μL of the SUMO-gp16 fusion protein was mixed with 200 μL wash buffer and 2.5 units of SUMO protease (Life Technologies), and the mixture was incubated at 4°C overnight. Ni-Sepharose 6 Fast Flow resin (GE Healthcare) was equilibrated with wash buffer and then incubated with the cleaved protein mixture for 60 min at 4°C. The resin containing the bound SUMO tag was pelleted in a microcentrifuge at 2,000 × *g* for 5 min, and the supernatant containing the native gp16 was recovered for use in biological assays.

The SUMO-gp16 plasmid was used as a template for creating gp16 mutants using the QuikChange Lightning site-directed mutagenesis kit from Agilent Technologies. Plasmid DNA was sequenced to verify the presence of the desired mutations.

### Molecular dynamics simulations

The initial Φ29 structure was taken from an atomic resolution cryo-EM reconstruction of a stalled motor during dsDNA packaging (PDB: 7JQQ) [1]. This pentameric structure consists of two unbound subunits and three subunit’s that are ATP-*γ*-S bound. Subunits that had no nucleotide in their binding pocket had ATP inserted, which was aligned based on distances to key residues as found in nucleotide bound subunits. Subunits that were already bound to ATP-*γ*-S had their sulfur replaced by an oxygen to create ATP-bound subunits. To create ADP-bound from ATP-bound subunits, their ATP’s gamma phosphate group was deleted. To create smaller composition varients, such as the dimers, unrequired parts were removed from the the newly modified fully ATP-bound pentamer. Mutated structures used in simulations were generated by replacing the side chains using LEaP, a package in AmberTools22 [24].

All-atom MD simulations were carrried out with the Amber22 package [24] using the ff19SB force field for protein interactions, OPC model for water [25], and bsc1 for dsDNA[26]. ATP and ADP parameters were taken from the Amber parameter database [27]. Hydrogen atoms connected to heavy atoms had their bonds constrained via the SHAKE algorithm. All proteins were simulated in a solvated truncated octahedral box with 14 Å of padding containing 150 mM of Cl^*−*^ and Na^+^ ions. Each system underwent energy minimization in the form of 600 steps of steepest descent and conjugate gradient and heating from 100 to 310 K over 200 ps. Each system was equilibrated in the isobaric-isothermal ensemble at 1 bar and 310 K. The equilibration period varied from system to system (around 100 ns for dimers) and was determined through the RMSD of the protein backbone of a single subunit. When the fluctuations of the system stabilized, we deemed the system as equilibrated and simulated for atleast 500 ns. All simulations were run on graphical processing units. Long-range interactions were corrected using the Ewald summation. Temperature and pressure control were implemented using the Monte Carlo barostat and the Langevin thermostat. For simulations to determine the minimal acceptable configuration to represent trans-interactions, no positional restraints were used. For the remaining dimer simulations (with and without dsDNA), weak positional restraints (2 kcal/mol/Å^2^) were used on the lid structure of the preceding subunit (residues 196-224) and the few residues where we expected the portal complex to contact the CTD for both subunits (residues 294, 297, 300, 301, and 329). CTDs were defined as residues 237 to 330 and NTDs as residues 3 to 192. Replicates were simulated using the same starting structure but different starting velocities determined from the Maxwell-Boltzmann distribution. These were compared for consistency using various metrics (RMSD, distances between key residues, and visualization of trajectory) and some comparisons are provided in the supporting information (SI Appendix, Fig. S5).

All structures and trajectories were visualized through ChimeraX [28] and VMD [29].

### Free energy calculations

The reaction coordinate was defined by the distance between the P_*γ*_ of the ATP bound to the reliant subunit and the center of mass of the two oxygens in the active group of the catalytic glutamate (E119) of the reliant subunit. The reaction coordinate was sampled using umbrella sampling by introducing harmonic restraints of varying spring constants, depending on the steepness of the energy landscape. Initially, umbrella sampling windows were set every 1 Å and a spring constant of 5 kcal/mol/Å^2^ was employed. Regions of the reaction coordinate that were not sampled sufficiently had additional windows introduced, with higher spring constants if needed, until proper coverage was achieved. Each window was sampled every 1 ps for at least 100 ns. The sampling was set up using the PLUMED library [30, 31]. The weighted histogram analysis method [32] was then used to calculate the free energy profile, technically known as the potential of mean force [33]. To calculate error bars of the free energy profile, the trajectory corresponding to each window was divided into four blocks, resulting in four psuedo independent free energy profiles. The error bar was the calculated as the standard error of the mean across the four profiles.

### Mutual information calculation

The MI of sidechains and backbone dihedral angles was calculated using CARDS [17] through the Enspara package [34] on 500 ns long MD simulations of the dimers, excluding the equilibration period. The holistic correlation between dihedrals X and Y is defined as 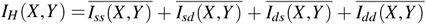, where the normalized mutual information between the structure of dihedral X and the structure of dihedral 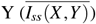, the dynamic state of X and the structure of Y (*I*_*ds*_(*X*,*Y*)), the structure of X and the dynamic of 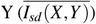, and the dynamical state of X and the dynamical state of 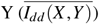 are added. The calculated MI had the outliers removed (determined by the interquartile range), was normalized, and then added to the PDB files as the b-factor for visualization. To verify for consistency, the MI of the first and second halves of the trajectory were compared, which we found to roughly converged.

### Structural relatives

Structures of Φ29 relatives that did not have existing structures (GA-1, SF5, and B103) were created by folding their subunit sequences into monomers using AlphaFold [21]. For each relative, two copies of the folded monomeric structure were superimposed onto a Φ29 ATP-ATP dimer (in the native cryo-EM conformation) to create their representative dimer structure. However, the relatives all share high sequence similarity to the Φ29 motor, so they are expected to form similar structures. The structure of Φ28 was taken from the Protein Databank (PDB: 7JQ6).

## Acknowledgments

This work was supported by NIH (Grant 2R01GM122979 to G.A., P.J.J, and M.C.M.). Computational resources were provided by the Duke Computing Cluster and by the ACCESS program supported by the National Science Foundation (Grants ACI-2138259, 2138286, 2138307, 2137603, and 2138296) [35]. Figure 7 was created using https://BioRender.com.

## Notes

### Competing Interest Statement

The authors have declared no competing interest.

