## Supplemental Information for "Regulation of Ordinal DNA Translocation Cycle in Bacteriophage Φ29 through Trans-Subunit Interactions"

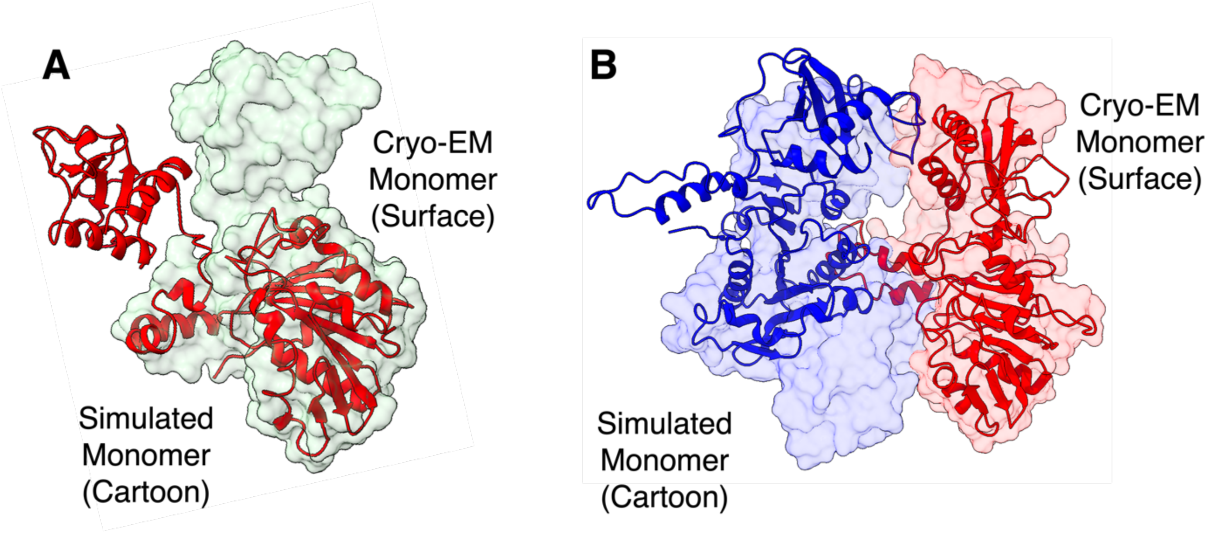


**Figure 1**. Structures obtained from monomer and dimer simulations. (A) Lack of support from adjacent subunits and dsDNA leads to conformations that would not be observed in the pentameric CryoEM structure. (B) Lack of DNA can cause drifting of the NTDs. The cartoon representations show a frame of the simulation, and the surface representation represents the conformation as seen in the CryoEM structure (PDB:7JQQ).

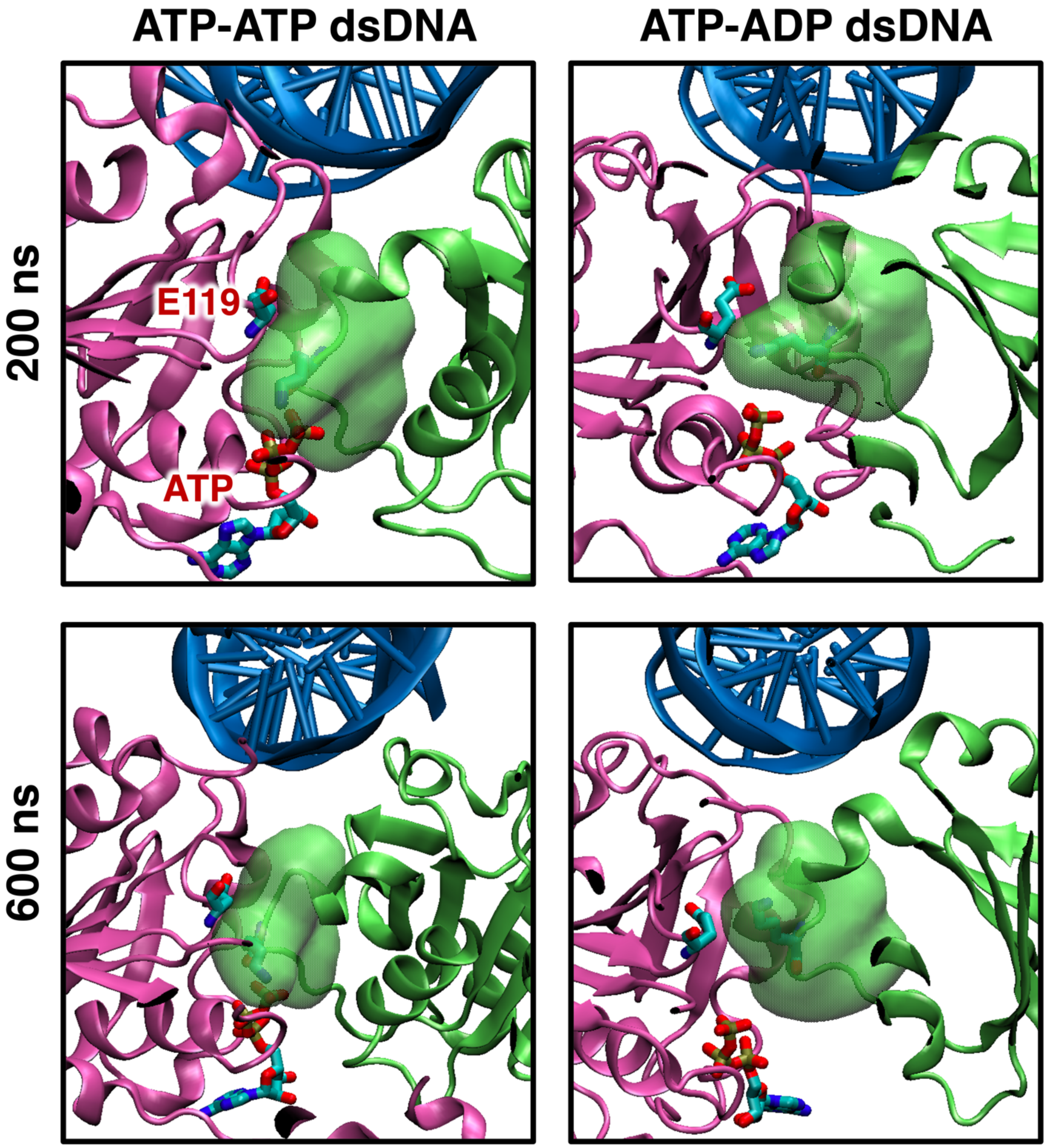


**Figure 2**. Dynamics of the obstructive motif. The position of the obstructive motif (green surface representation) of the preceding subunit (green cartoon representation) relative to the ATP and catalytic glutamate (E119) of the reliant subunit (mauve cartoon representation). The left column contains frames from the simulated ATP-ATP DNA dimer and right column from of the ATP-ADP DNA dimer. At 200 ns of the simulation (top row), both ATP-ATP and ATP-ADP dimers have the obstructive motif positioned between the catalytic glutamate and ATP. At 600 ns (bottom row), the obstructive motif remains in a similar position in the ATP-ATP dimer, but has moved out of the way in the ATP-ADP dimer. DNA is in the blue cartoon representation.

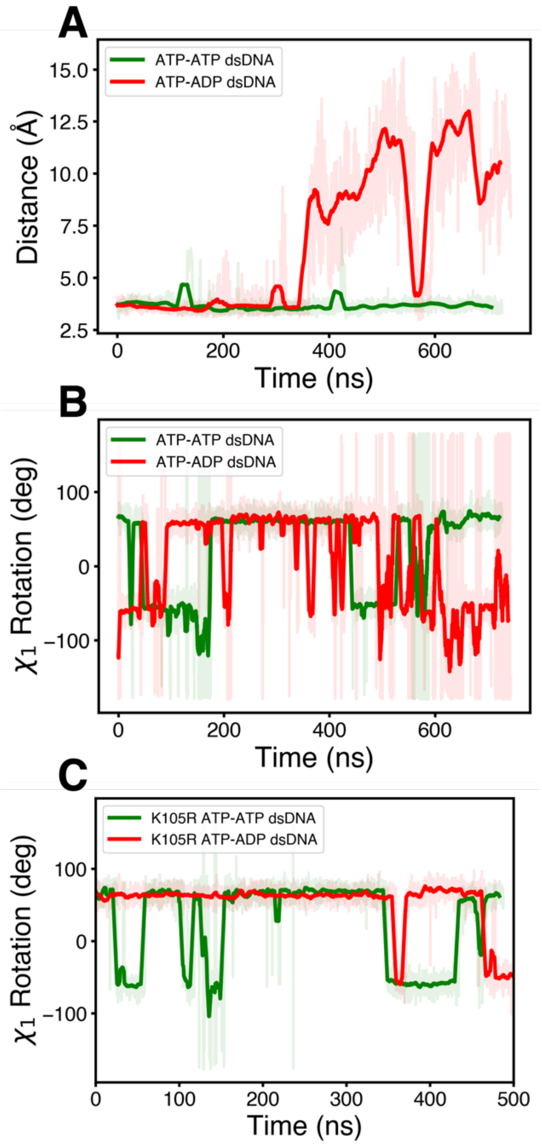


**Figure 3**. Movement and rotation of the preceding subunit's obstructive lysine (K105). (A) Distance between the P\textsubscript{$\gamma$} of reliant subunit's ATP and preceding subunit's K105. The distance remains small in an ATP-ATP DNA dimer (green) but can increase in the ATP-ADP DNA dimer (red). (B) Rotation of the $\chi_1$ angle of K105. The $\chi_1$ dihedral is more stable in the ATP-ATP dimer compared to the ATP-ADP dimer, seems to be correlated to the increased separation seen in A. (C) Rotation of the $\chi_1$ angle of K105 when mutated into an arginine (K105R). The rotation of the $\chi_1$ angle is decreased in the mutated ATP-ADP dimer when compared to the native dimer. Shaded regions represents the value at every frame (recorded at 10 ps intervals) and the solid lines are the moving averages (every 300 frames for panel A, every 50 frames for panel B and C).

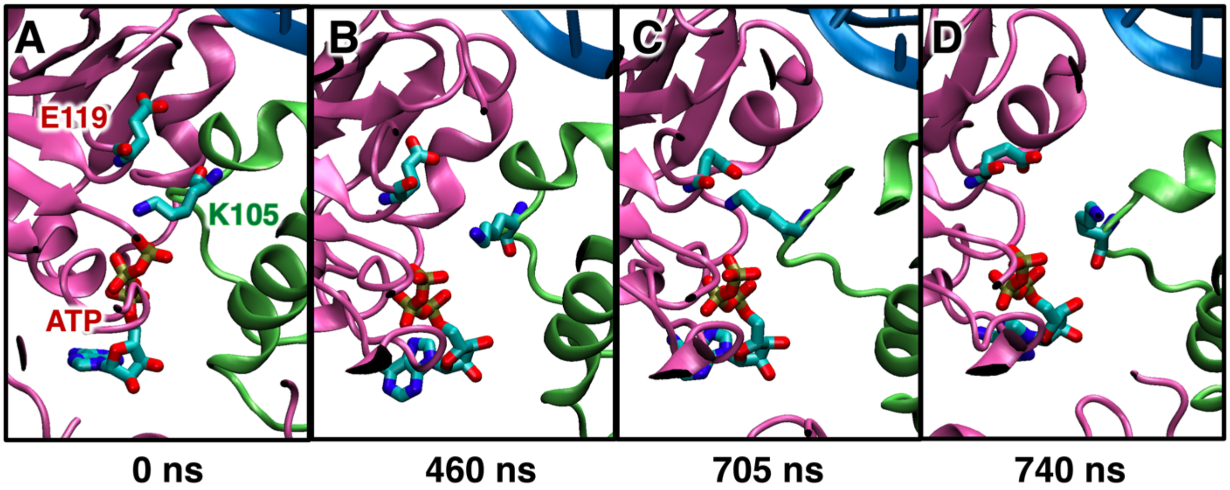


**Figure 4**. Active site of an ATP-ADP DNA dimer, showcasing the ATP and catalytic glutamate (E119) of the reliant subunit (mauve cartoon) and the obstructive lysine (K105) of the preceding subunit (green cartoon). The method by which the obstruction and K105 moves in and out of position appears to be mostly dictated by the rotation of the side chain. At 0 ns, it is interacting with the gamma phosphate of ATP (A), at 460 ns it rotates out of the way (B), at 705 ns it is seen interacting with E119 (C), and at 740 ns it is interacting with neither (D). DNA can be seen in the top right of the frames (blue cartoon).

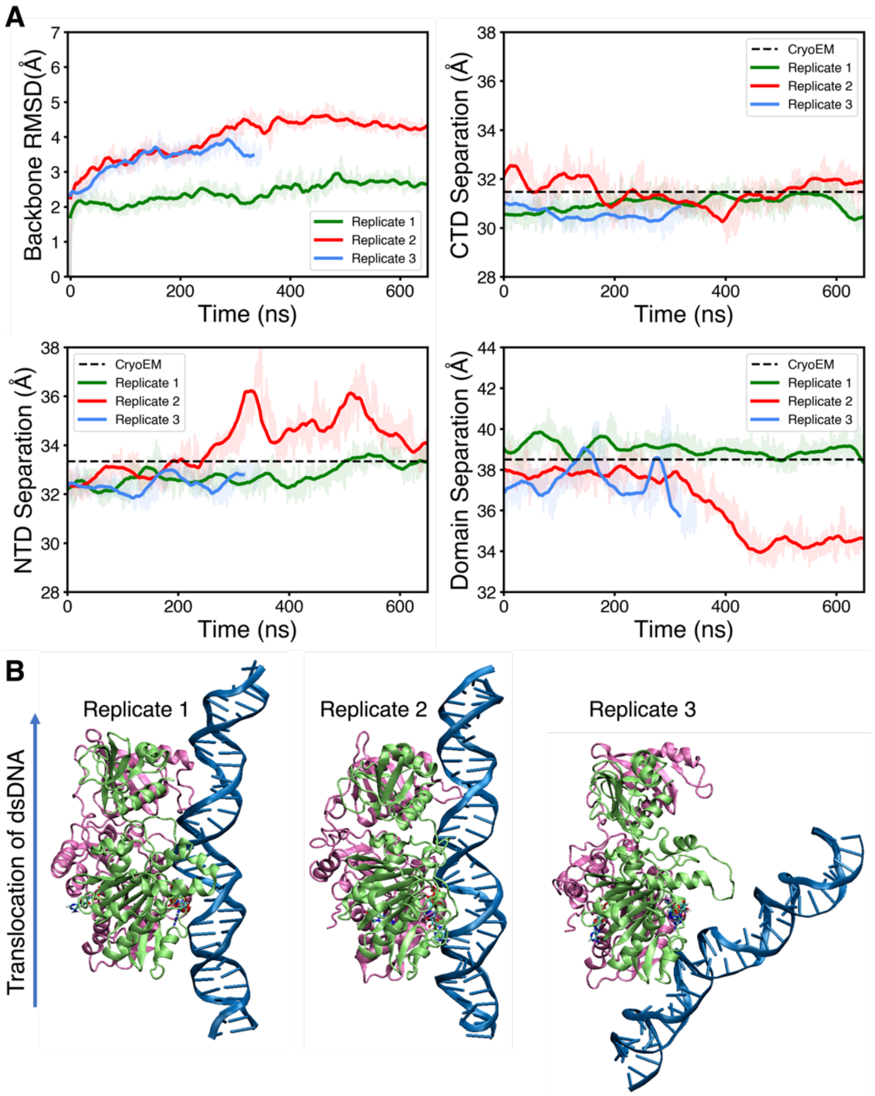


**Figure 5**. Comparisons of three ATP-ATP dsDNA replicate simulations. These were initiated from the same starting structure, but with different starting velocities. (A) Protein backbone RMSD in reference to each replicate's own starting frame (top left). The distances between the center of masses of the CTDs of the two subunits (top right), distances between the center of masses of the NTDs of the two subunit (bottom right), and distances between the center of masses of the CTD and NTD of the reliant subunit (bottom left). Even though a large variation in the RMSD can be seen, the domain distances were consistent with each other and the CryoEM structure. Shaded regions represents the value at every frame (recorded at 10 ps intervals) and the solid lines are the moving average (every 150 frames for RMSD panel, every 300 frames for the remaining panels). (B) Final frames of each replicate (green is preceding subunit, mauve is reliant subunit, and blue is DNA). Dissociation of the DNA and subunits can be seen in replicate 3, and thus its simulation was ended early (right).

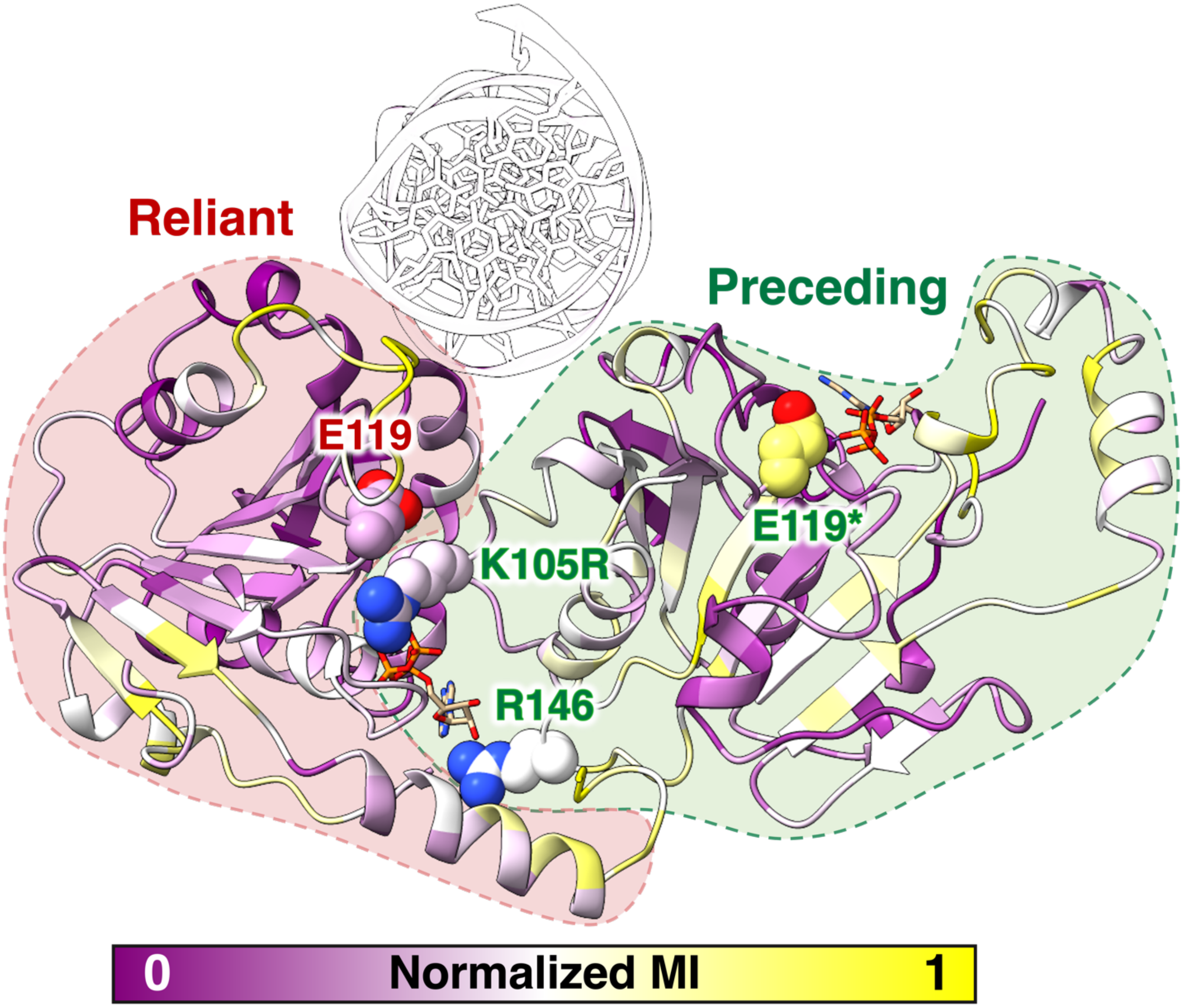


**Figure 6**. Mutual Information (MI) calculations of an ATP-ATP dsDNA dimer which contains a K105R mutant in the preceding subunit. The target residue is the preceding subunit's catalytic glutamate (E119) and the correlation from E119 to K105R is not strong.}
